# Transposable Element Exprssion in Tumors is Associated with Immune Infiltration and Increased Antigenicity

**DOI:** 10.1101/388215

**Authors:** Yu Kong, Chris Rose, Ashley A. Cass, Martine Darwish, Steve Lianoglou, Pete M. Haverty, Ann-Jay Tong, Craig Blanchette, Ira Mellman, Richard Bourgon, John Greally, Suchit Jhunjhunwala, Matthew L. Albert, Haiyin Chen-Harris

## Abstract

Profound loss of DNA methylation is a well-recognized hallmark of cancer. Given its role in silencing transposable elements (TEs), we hypothesized that extensive TE expression occurs in tumors with highly demethylated DNA. We developed *REdiscoverTE,* a computational method for quantifying genome-wide TE expression in RNA sequencing data. Using The Cancer Genome Atlas database, we observed increased expression of over 400 TE subfamilies, of which 262 appeared to result from a proximal loss of DNA methylation. The most recurrent TEs were among the evolutionarily youngest in the genome, predominantly expressed from intergenic loci, and associated with antiviral or DNA damage responses. Treatment of glioblastoma cells with a demethylation agent resulted in both increased TE expression and de novo presentation of TE-derived peptides on MHC class I molecules. Therapeutic reactivation of tumor-specific TEs may synergize with immunotherapy by inducing both inflammation and the display of potentially immunogenic neoantigens.

**One Sentence Summary:** Transposable element expression in tumors is associated with increased immune response and provides tumor-associated antigens

## Main Text

One of the key insights from the Human Genome Project was the discovery that at least 50% of the human genome is comprised of repetitive sequences, most of them derived from transposable elements (TE) of unknown function (*1*). Like all eukaryotic genomes (*2–4*), the human genome encodes multiple defense strategies to silence the expression and mobility of TEs, including epigenetic repression by DNA methylation. In the context of cancer pathogenesis, transformation to a malignant state is frequently accompanied by a global loss of DNA methylation (*5, 6*). Conceivably, this global epigenetic dysregulation in cancer cells leads to extensive TE expression, which in turn impacts both tumor-cell intrinsic functions and anti-tumor immune response. Indeed, there is converging evidence from *in vitro* and *in vivo* preclinical studies that treatment with DNA demethylation agents triggers expression of certain human endogenous retroviruses (HERV) in cancer cells and activate a robust type I interferon response (*7, 8*). As such, epigenetic therapies may have the potential to sensitize patients to immune checkpoint therapy by inducing inflammation (*9*) and by the formation of immunogenic TE-derived peptide-MHC class I complexes. In support of this possibility, in a hematopoietic stem cell transplantation trial in kidney clear cell carcinoma, tumor regression was associated with donor T cells’ recognition of HERV-E antigens (*10, 11*).

A major impediment to understanding TE expression and its potential relevance to tumor immunity is the analytic challenge of accurate quantification of short-read sequences from repetitive regions in the transcriptome. Standard pipelines typically discard repetitive reads (*12*). Most current TE quantification techniques rely on: mapping reads to consensus sequences, exclusion of multi-mapping reads, or step-wise operations that could introduce considerable biases (*13–16*). We have developed and benchmarked a new method, *REdiscoverTE,* to quantify TE expression from RNA-seq data, then applied it to thousands of cancer and normal tissue samples from The Cancer Genome Atlas (TCGA) and the Cancer Genome Project (CGP) (*17*).

In this study, we characterized the patterns of genome-wide TE expression, related expression to DNA methylation, and identified numerous ways in which TE expression may impact innate and adaptive immune responses to the tumor.

## Results

### Genome-wide TE expression quantification approach

*REdiscoverTE,* whose features are detailed in the Methods, was devised to allow simultaneous quantification of all annotated genes and TEs expressed in the human genome using a comprehensive human transcriptome reference that included all human RepeatMasker sequences (*18*) (**Fig. 1A, Table S1**). To mitigate the uncertainty associated with mapping reads to repetitive features, we leveraged a recently developed lightweight mapping approach for isoform quantification, *Salmon,* which uses an expectation-maximization (EM) algorithm to assign multi-mapping reads probabilistically to transcripts, based on evidence from uniquely mapped reads **(19)**.

**Fig. 1.**
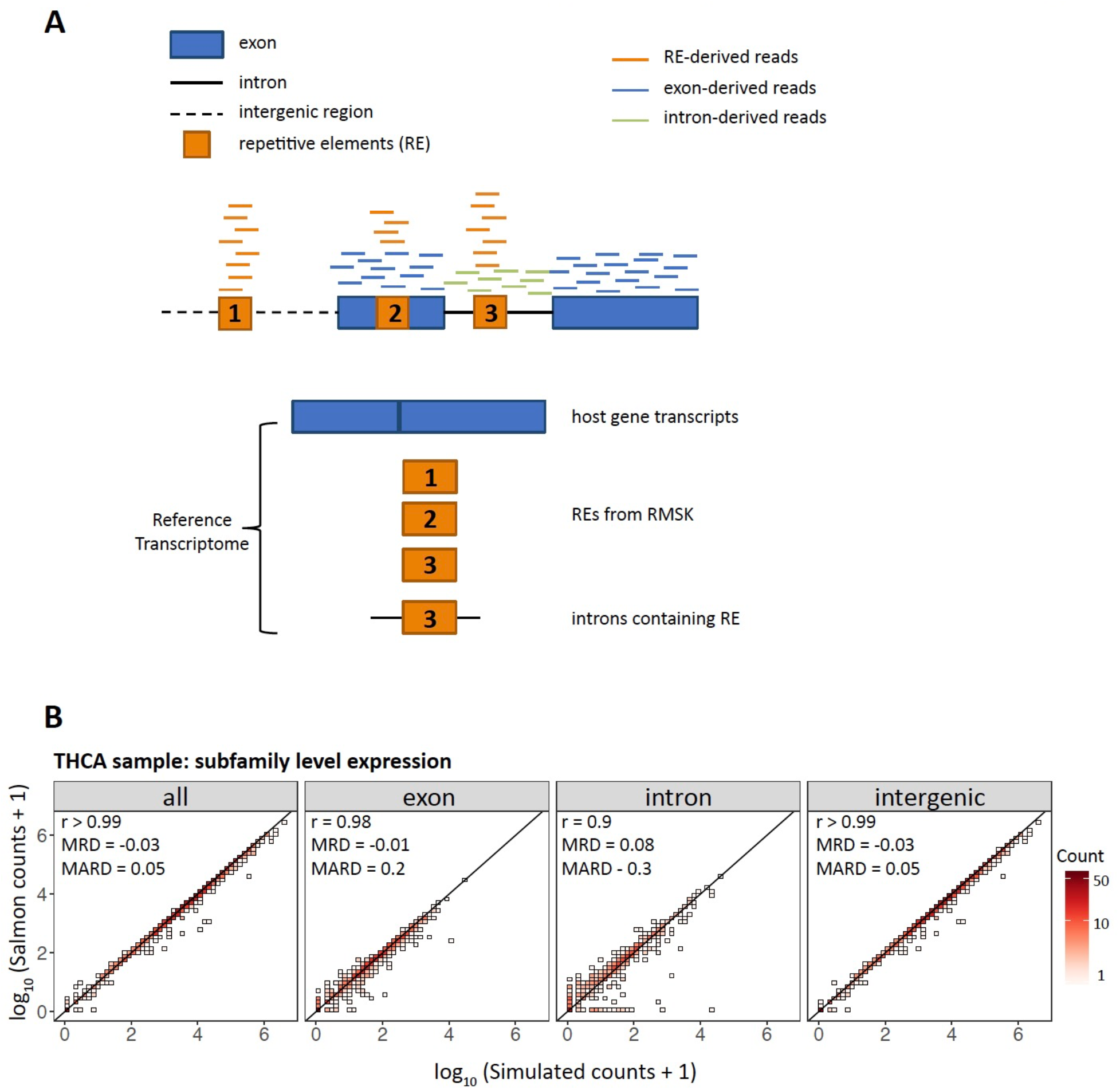
*REdiscoverTE* reference transcriptome and performance benchmarking. A. REdiscoverTE whole transcriptome reference for RE quantification. Schematic depicts short reads mapping to host gene features (exons, introns) and REs either embedded within host genes or intergenic regions. RE genomic locations are derived from RepeatMasker (RMSK). Reads stemming from repetitive elements (REs), exons, and introns are illustrated in orange, blue and green, respectively. B. Benchmarking the accuracy of RE quantification by REdiscoverTE with simulation. Two-dimensional histogram comparing REdiscoverTE quantification to simulated RE expression generated based on a TCGA THCA sample. Expression is aggregated to the subfamily level. Left to right: all RE expression regardless of genomic context, exonic RE expression, intronic RE expression, intergenic RE expression. Performance accuracy is measured in terms of Spearman correlation coefficient (r), mean relative difference (MRD), mean absolute relative difference (MARD).

While the *REdiscoverTE* transcriptome reference includes all 5 million human RepeatMasker repetitive elements, we restricted our downstream analysis to TEs only, which encompass 4 million elements classified into 1,052 distinct TE subfamilies in 5 classes: long interspersed nuclear elements (LINE), short interspersed nuclear elements (SINE), long terminal repeats (LTR), SINE-VNTR-Alu (SVA), and DNA transposons (**Fig. S1A**). We were primarily interested in measuring the total transcriptional output from a group of related TEs regardless of their genomic location. Therefore, after quantification at the individual locus specific-level, we chose to aggregate expression for individual elements to the TE subfamily level. We further divided the aggregate expression for each TE subfamily into exonic, intronic and intergenic expression by stratifying all elements under a given TE subfamily by their genomic locations with respect to host genes as defined in Gencode (*20*). For example, of the 1,610 annotated instances of L1HS, 951 are located in the intergenic regions, 654 in host gene intronic regions and 5 in host gene exons. Here, L1HS intergenic expression was defined as the aggregate expression from the 951 elements within the intergenic regions. Other subfamilies were treated similarly.

We first benchmarked the performance of *REdiscoverTE* with extensive simulations using *RSEM (21*) and found *REdiscoverTE* to be highly accurate (**Fig. 1B, Fig. S1C-G**). While *REdiscoverTE* can accurately quantify expression in a locus-specific manner, we found the approach of count aggregation of expression to the subfamily level achieved much higher the accuracy of TE quantification, likely by reducing mapping noise observed at the individual element level (**Fig. S1F-G**, more details in **Supplementary Notes**). *REdiscoverTE* performed best on intergenic TEs, followed by exonic, then intronic TEs (**Fig. S1F-G**). Exonic TE expression, which comprised a minority of the total TE read fraction, was excluded from further analysis to rule out the potential confounding expression from overlapping host genes and ease down-stream interpretation.

For further confirmation, we performed a direct comparison of *REdiscoverTE* to a previously published study on a restricted set of 66 human endogenous retroviruses (HERVs) in TCGA by Rooney et al. (*15*). We found the results were generally consistent (median r = 0.76, **Fig. S1K**), particularly for 3 previously identified tumor-specific HERVs: ERVH-5, ERVH48-1 and ERVE-4 (**Fig. S1J**). However, *REdiscoverTE* has the advantage of efficiently capturing expression by all REs.

Finally, we compared *REdiscoverTE* to *Repenrich(14*), a two-step alignment-based TE expression quantification method that first aligns reads to host genes using Bowtie (*22*), then quantifies TE expression. Benchmarking against simulated data, we showed that *REdiscoverTE* performed with higher accuracy and computational efficiency (**Fig. S1H-I**), while *RepEnrich* showed significant over-estimation of TE expression. Likely reasons for RepEnrich’s overestimation include the addition of padding sequences in its TE reference and the assignment of reads that belong to genes and retained introns to overlapping TEs. *REdiscoverTE* overcomes these biases with the inclusion of intron sequences in its transcriptome and simultaneous host gene/RE quantification.

### TE expression is dysregulated in cancer

To characterize the landscape of TE expression in cancer, we applied *REdiscoverTE* to 7,345 TCGA RNA-seq samples (containing 1,232 tumor and matched normal samples, the rest are tumor samples without normals) across 25 cancer types. For validation of our findings in select cancer types, we also analyzed an additional 405 tumor and matched normal RNA-seq samples across 5 cancer types (**Table S2**) from CGP (*17*). Both datasets were generated from poly-dT RNA-seq library preparations, which can capture poly-adenylated TEs transcripts. On average, 1% of RNA-seq output mapped to TEs (**Fig. S2A**). Notably, TE expression was observed from all TE classes (N=5) and most families (N=43), including both retrotransposons and DNA transposons (**Fig. S2B**). For each TE class, the bulk of expression stem from intergenic regions (**Fig. S2C**), suggesting autonomous TE expression from intergenic loci compared to read-through transcription in host, protein-coding genes. Human DNA transposons had been thought to be completely inactive, based on the lack of evidence for transposition in the human genome(*23*). Although these data cannot address transposition, our results, coupled with other recent studies (*24, 25*), suggest active gene expression by DNA transposons.

In all cancer types, TE expression was detected in both tumor and matched normal tissues, suggesting basal levels of TE transcriptional activities in normal tissues. Across the two datasets, 10 cancer types showed a significantly higher proportion of reads mapping to TEs in tumor compared to matched normal tissues, suggesting that TE expression may be particularly active in these cancers; the reverse was observed in 4 cancer types (**Fig. S2A, Fig. S2D, Fig. S2E**). Consistent with this, differential expression analysis of tumor samples with respect to matched normals (*26–28*) revealed that stomach, bladder, liver, and head and neck tumors show predominantly over-expression of TEs, while thyroid, breast, kidney chromophobe and lung adenocarcinoma tumors show predominantly reduced TE expression compared to normal (**Fig. 2A**). Across all tumor types, many TEs showed differential expression: out of 1,052 TE subfamilies, 587 were differentially expressed in at least one TCGA cancer type studied, of which, 463 are over-expressed in at least one cancer type (**Fig. 2A-C, Table S3, S4**). Notably, for many cancer types, a number of TE subfamilies showed low expression in normal samples but significantly higher expression in tumor samples (**Fig. 2B**). The TE class LTR showed the highest number of over-expressed subfamilies followed by DNA and LINE. However, this enrichment pattern was largely consistent with the total number of subfamilies defined in these TE classes, hence there is little evidence of class preference for over-expression (**Fig. 2C**). The pattern of over-expression in tumor versus normal tissue was highly consistent across the two data sets (TCGA and CGP), suggesting that the TE expression profile may be characteristic of tumor/tissue type (**Fig. 2D**).

**Fig. 2.**
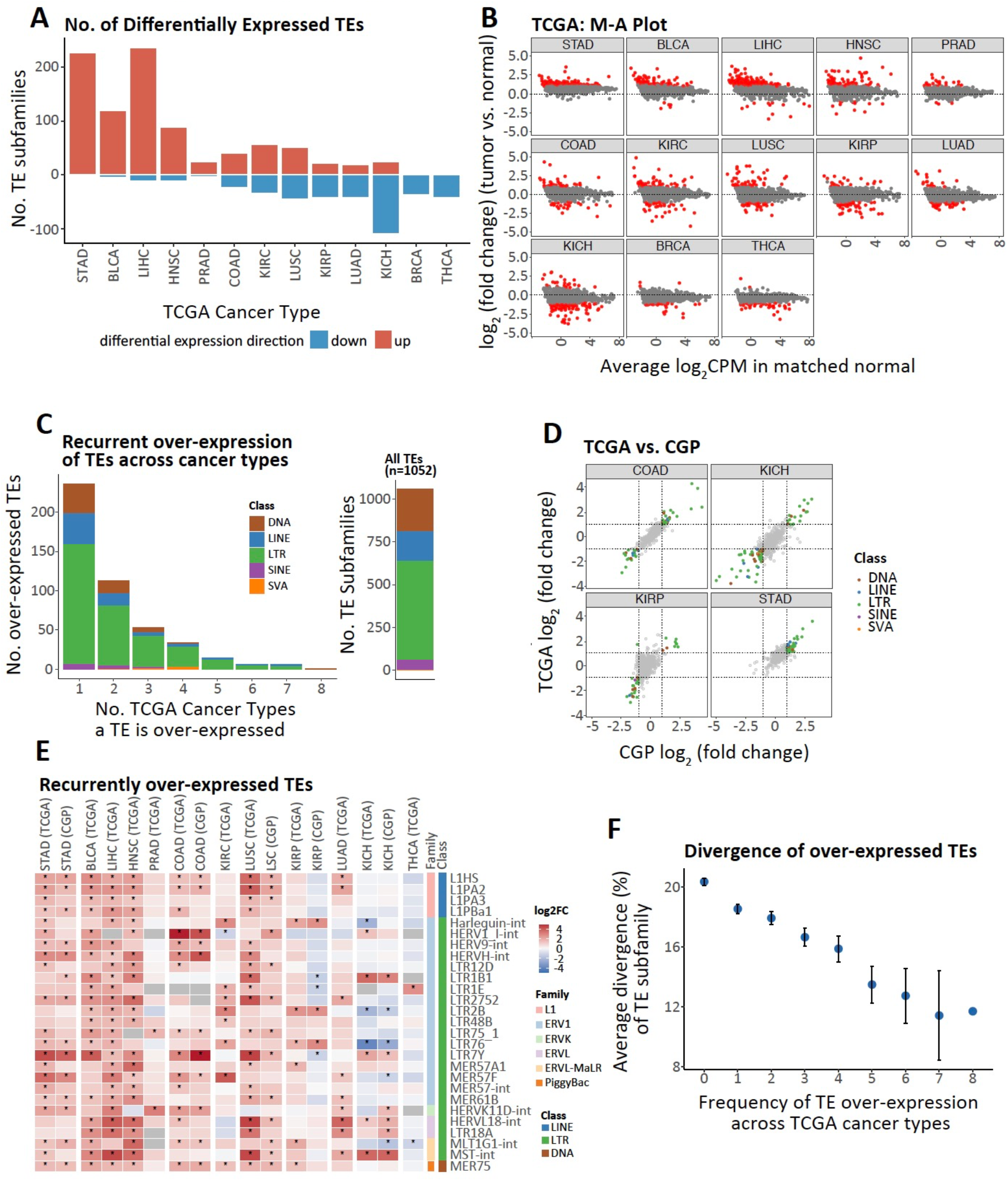
TE expression is dysregulated in cancer. (All differential expression analysis are based on intergenic TE expression and performed on matched tumor-normal sample pairs) A. Number of differentially expressed TE subfamilies in 13 TCGA cancer types. Red bars: number of significantly over-expressed TE subfamilies. Blue bars: number of significantly under-expressed TE subfamilies. Cancer types are ordered from left to right by the ratio between number of over-expressed and under-expressed TE subfamilies. Significance is defined at log2 fold-change (FC) > 1 and FDR < 0.05. B. M-A plot showing TE expression FC of tumor over normal as a function of mean normal tissue TE expression (log2 counts per million; CPM) for 13 TCGA cancer types. Each point is one TE subfamily. Red: significantly differentially expressed TE subfamilies. Cancer types are ordered as in Fig. 2A. C. Left: histogram of TE subfamilies by number of TCGA cancer types in which they are over-expressed to show recurrence of over-expression. In total, twenty-seven TE subfamilies were over-expressed in at least 5 cancer types. Right: number of TE subfamilies in each of 5 TE classes as defined by Repeatmasker GRCh38. D. Comparison of TE differential expression profile (tumor vs. matched normal) between TCGA and CGP RNA-seq data on matching cancer types. E. FC of expression for the 27 TE subfamilies are selected based on Fig. 2C. Heatmap colors indicate log2 FC (tumor vs. matched normal) values; columns are ordered as in Fig. 2A. CGP data are grouped with corresponding TCGA cancer types. F. Younger TEs tend to be more frequently over-expressed in multiple tumor types. Average divergence of TE (proxy for age) was calculated for each TE subfamily. The plot summarizes the average divergence for TE subfamilies at a given frequency of TE over-expression across cancer types (grouping of TEs are based on Fig. 2C). Error bars indicate standard error of the mean of average sequence divergence of TE subfamilies at a given prevalence.

A minority of TEs were recurrently expressed in multiple cancer types: 61 TE subfamilies were significantly over-expressed in at least 4 cancer types (**Fig. 2C**). Among these, L1HS, the human specific subfamily of the LINE1 family, was over-expressed in 8 cancer types (**Fig. 2E, Fig. S2G**). A number of TE subfamilies belonging to the human endogenous retrovirus (HERV) group were also consistently over-expressed. Full-length HERV sequences are characterized by the presence of two LTRs flanking a proviral genome (*23, 29*). HERVK11D-int, a member of the HERVK family (HERVK is the youngest family of HERVs), was over-expressed in 7 cancer types (**Fig. 2E, Fig. S2G**). LTR7Y, the youngest variant of the LTRs associated with HERVH (*30*), was over-expressed in 8 types of tumors. We discovered that coincident with the over-expression of LTR7Y, HERVH-int, the proviral portion of HERVH, was over-expression in six of the same cancer types (**Fig. 2E, Fig. S2H, Fig. S2I, Table S4**), suggesting either shared regulation with LTR7Y or potential expression of full length HERVH sequence. A similar pattern of co-expression was also found for HERVL18-int and its associated LTR element LTR18A, and to some extent L1HS and SVA_F (**Fig. S2I**).

The evolutionary age of TE subfamilies can be estimated by their divergence from their consensus sequence (Smit, 2013). By this method, we found that the younger TE members are generally more consistently over-expressed across tumor types, in keeping with the above observation of prevalent over-expression of L1HS, HERVK and HERVH, which are known to be among the youngest TEs in the human genome (**Fig. 2F, Fig. S2F**). This suggests that younger TEs may be more likely to be active in tumor genomes, due either to more intact sequences and thus preserved promotors and transcriptional potential or to fewer overlapping mechanisms of silencing.

### Reactivation of TEs is associated with loss of DNA methylation

To elucidate the role of DNA methylation alterations in TE expression, we examined TCGA DNA methylation changes from normal to tumor at both the global and TE-proximal level using TCGA Illumina 450K array data, which captured 70K CpG sites overlapping with 1,007 TE subfamilies (**Fig. S3A**). Across 10 cancer types, we observed that, much like differential TE expression, the global pattern of differentially methylated CpGs (DMCs) was cancer type-dependent: CpGs in certain tumor types are often highly demethylated, e.g. LIHC and HNSC, while in other types they are often over-methylated, e.g. KIRP and PRAD (**Fig. 3A, Fig. S3B, Fig. S3C, Table S5**). Furthermore, in all cancer types considered, we discovered a strong enrichment of demethylation at CpG sites located within TEs, as compared to background demethylation at all CpG sites on the 450K array (**Fig. 3A, Fig. S3C, Table S5**). More strikingly, in all cancer types other than KIRP, tumor samples showed dramatic enrichment of de-methylation relative to over-methylation at TE-proximal CpG sites, indicating that loss of DNA methylation in these specific regions may be a common tumor pathology. Consistent with the known role of DNA methylation for TE silencing, the extent of demethylation, in terms of log-ratio of demethylated vs. over-methylated TE CpGs, was strongly associated with the extent of TE over-expression across cancer types (**Fig. 3B**).

**Fig. 3.**
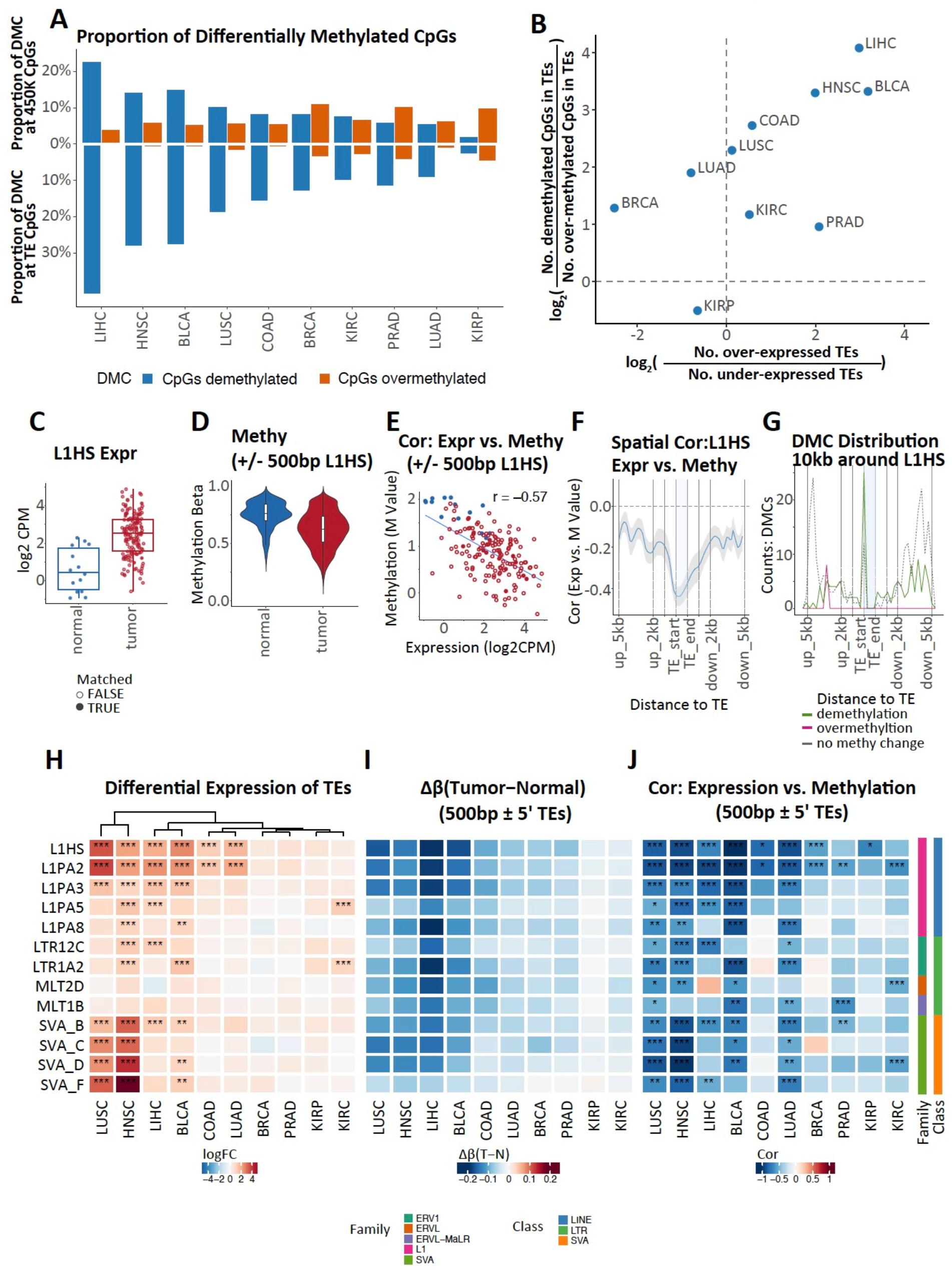
TE expression in cancer is associated with global and proximal epigenetic dysregulation. A. Global differential methylation states across TCGA cancer types. Criteria for significant differentially methylated cytosines (DMCs): absolute (delta beta value) >= 10%, FDR < 0. 05. Top: proportion of DMCs among all Illumina 450K CpG sites. Bottom: proportion of DMCs at CpGs within TEs. Blue: proportion of de-methylated DMCs among all CpG sites. Orange: proportion of over-methylated DMCs among all CpG sites. B. The extent of TE mRNA overexpression is strongly correlated with the extent of CpG demethylation within TEs. Each point represents one cancer type. Horizontal axis: log2 ratio between the number of over-expressed TE subfamilies and the number of underexpressed TE subfamilies. Vertical axis: log2 ratio between the number of de-methylated DMCs in TEs and the number of over-methylated DMCs in TEs. C-G Association between L1HS intergenic expression and its DNA methylation state in BLCA. Analysis was conducted using all samples. C. L1HS intergenic expression in normal and tumor samples. Blue: normal sample. Red: tumor samples. Filled circle: tumor samples with matched normal. Open circle: tumor samples without matched normal). D. L1HS proximal CpG M-value in normal and tumor samples. Blue: normal samples. Red: tumor samples. CpG sites are from 500bp+/− regions around intergenic L1HS 5’ bp (taken as TSS). E. Correlation between intergenic L1HS expression and methylation M value F. Spatial correlation between L1HS expression and CpG methylation M value 5kb+/− L1HS. Correlation was calculated for all samples at each CpG site, then smoothed with binsize = 500bp. Shading indicates 95% confidence interval. G. Spatial distribution of de-methylated CpG (green), over-methylated CpGs (red) and CpGs with no methylation change (grey, dashed) 5kb+/− around L1HS. Binsize = 500bp. H-J. Examples of TE subfamilies with significant negative correlation (cor<= −0.4 & FDR<0.05) between intergenic expression and methylation in more than 4 types of tumors. Results are based on matched tumor-normal samples. H. Tumor vs. Normal differential expression of select TE subfamilies. Heatmap colors: log2 FC. Significance level *: logFC>1&FDR<0.05; **: logFC>1&FDR<0.01; ***: logF C> 1&FDR<0.001. I. Tumor – normal average delta beta value in 500bp+/− regions around 5’ bp of all intergenic loci of given TE subfamily. J. Correlation between intergenic TE expression and methylation M values around 500bp +/− 5’ bp of intergenic TE. Heatmap colors: correlation (cor) coefficient. Significance level *: abs(cor)>=0.4&FDR<0.05, **: abs(cor)>=0.4&FDR<0.01; ***: abs(cor)>=0.4&FDR<0.001

To gain better resolution on methylation patterns around TEs, we performed sample-level correlation and spatial analysis of DMCs. We illustrate the approach with intergenic L1HS expression and methylation analysis in BLCA. Relative to normal bladder tissue, L1HS is significantly over-expressed in bladder tumor (log2FC = 2.3, p=4°10^-7^, limma, **Fig. 3C**); and an inverse relationship is seen for methylation marks, with L1HS proximal CpG sites being significantly demethylated in tumor compared to normal tissue (p=2°10^-16^, two sided t-test, **Fig. 3D**). Across samples, the average L1HS methylation level was significantly inversely correlated with aggregate L1HS expression level (p=2°10^-15^, cor = −0.57, **Fig. 3E**). Next we created a spatial correlation profile between methylation level and aggregate expression level for a 10kb region around all L1HS loci and observed a deepening inverse correlation at the 5’ end of L1HS in BLCA (**Fig. 3F**). Finally, a DMC spatial enrichment profile for the same 10kb region further confirmed a strong enrichment of demethylated DMCs at the 5’ end of L1HS in BLCA (**Fig. 3G**). These results together establish that intergenic L1HS activity in tissue is influenced by the DNA methylation state at its 5’ end.

We extended this correlation analysis to 1,007 TE subfamilies in 10 cancer types, and discovered a strong inverse correlation for 431 TE subfamilies (**Fig. S3D, Table S6**), 262 of which showed significant over-expression in at least one cancer type. We highlight 13 TEs subfamilies from the LINE, LTR and SVA class that showed recurrent significant inverse correlation between expression and proximal DNA methylation across cancer types (**Fig. 3H, Fig. 3I, Fig. 3J**). Of note, we found that the over-expression of SVA, the youngest active group of retroelements in hominids (*31*), is strongly associated with proximal DNA de-methylation, particularly in head and neck squamous cell and lung squamous cell carcinoma (**Fig. 3J, Fig. S3F**).

As noted above, we observed predominantly reduced levels of TE expression in tumor compared to normal tissue in a subset of cancer types (thyroid, breast, kidney chromophobe and lung adenocarcinoma). We examined DNA methylation status at 6 recurrently down-regulated TEs, but found no clear association between methylation and TE expression (**Fig. S3D**).

Together, these data demonstrate that the over-expression of many TEs in tumor is associated with loss of DNA methylation, particularly at TE-proximal CpG sites, suggesting that a major mechanism of TE reactivation may be targeted loss of DNA methylation near TEs.

### TE expression is associated with DNA damage and immune response

We next tested the hypothesis that tumor TE expression can impact cellular and immune response within the tumor by examining its relationship to transcriptional activities of major cellular pathways. Twenty-four pathways were considered, including 8 related to cancer (e.g. P53 signaling), 6 related to DNA damage response (e.g. homologous recombination) and 8 related to immune response (e.g. type I IFN response) (*32*) (**Table S7**). For each pathway of interest, we first scored its overall activity in the tumor samples using singular-value weighted gene expression of the associated gene set. Scoring bulk tissue for expression signatures creates an interpretation challenge: as both tumor and non-tumor cells contribute to the mRNA signal, any observed differences across samples may result from differential tumor cell TE (or other gene) expression *or* from differences in sample cellularity, i.e., in the relative abundance of tumor, immune, or stromal cell content. To isolate the contribution of TE to variable expression of these pathways, we compared two statistical models: (*1*) a cellularity-only model that relates pathway scores to estimates of sample tumor purity plus lymphoid and myeloid cell content (*33*) (**Fig. S4A, Fig. S4B**); and (*2*) a full lasso regularized regression model (*34*) that relates pathway scores to the expression of 1,052 TE subfamilies in addition to the 3 tumor cellular components (tumor, lymphoid, myeloid). The difference between the goodness of fit (the r-squared values) of the two models was interpreted as the total explanatory power from all TE expression to each pathway, since the two models both account for cellularity variations in the samples. As Lasso is a statistical technique that selects representatives of correlated variables, the models also helped to identify representative top TE contributors to the variable expression of each pathway in question (**Table S8**).

Comparing the full lasso model with the cellularity model revealed that total TE expression accounted for substantial explanatory power to expression of many pathways with change in r-squared values exceeding 0.4, including DDR, type I IFN response, antigen processing pathways, cell cycle, P53 signaling and epithelial-mesenchymal transition (**Fig. 4A, Fig. S4C, Table S8**). Specifically for DDR pathways, the mean r-squared-values across cancer type was substantially higher in the full lasso model (0.74±0.14) than the cellularity only model (0.18±0.10). MER75 (piggyBac family, DNA transposon), MER4A (ERV1 family), MER54A (ERV3), MER67A (ERV1) were identified by the full lasso model as top predictors for DDR activities (**Fig. 4B, Fig. S4D, Fig. S4E**).

**Fig. 4.**
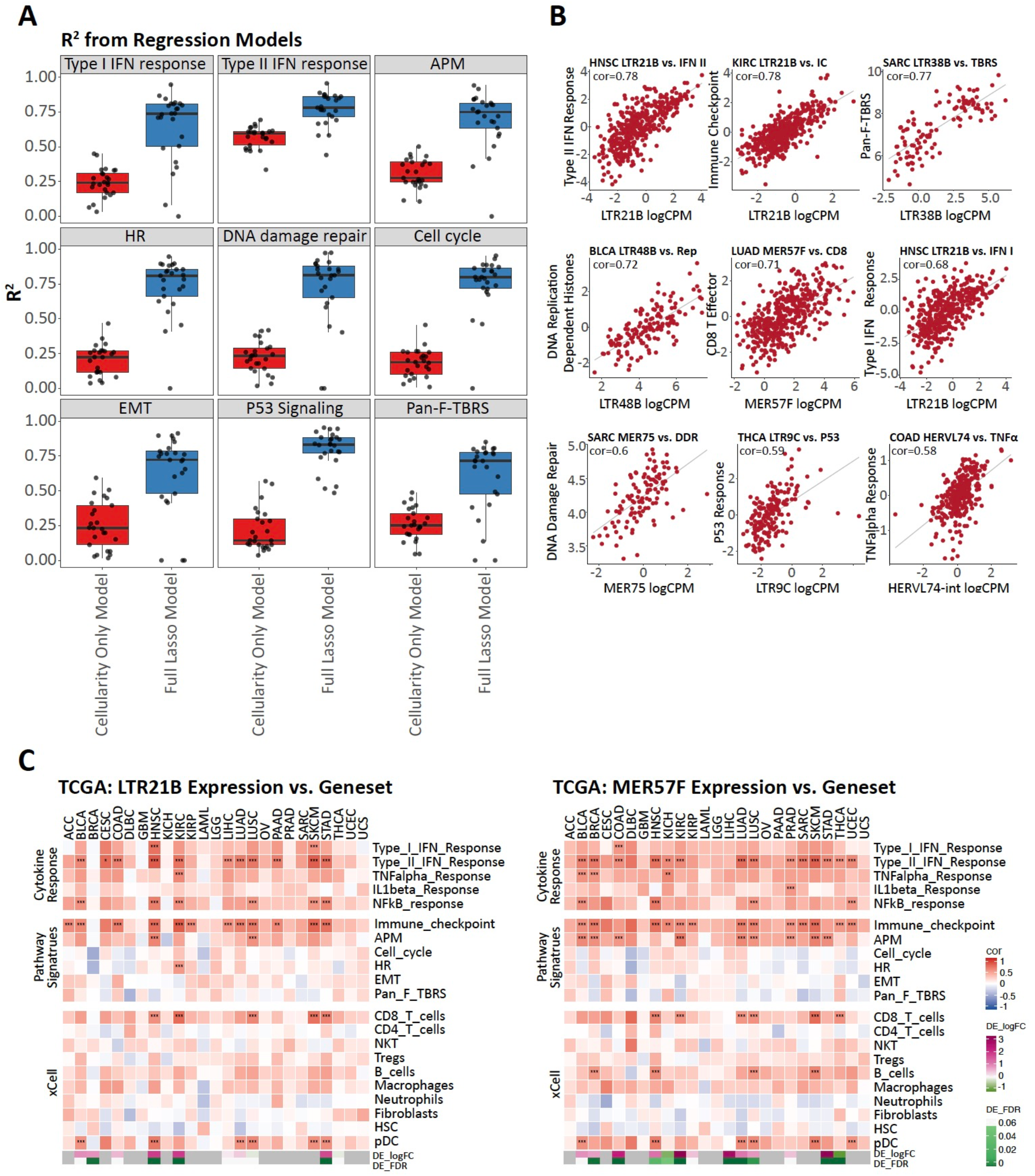
TE activity is associated with cellular and immune response in the tumor. A. R2 estimated by two models for each of 9 gene signature scores. Each panel is one gene signature, each point is one cancer type. HR: homologous recombination. APM: antigen processing machinery. EMT: epithelia-mesenchymal-transition. Pan-F-TBRS: pan-fibroblast TGFbeta response signature. Red: R2 estimated by linear model to predict gene signature scores using 3 covariates -- tumor content, lymphoid and myeloid, as predictors. Blue: R2 estimated by Lasso model taking 3 covariates and expression level of all 1,052 TEs as predictors B. Examples of positive correlations between gene signature scores and TE expression levels in different TCGA cancer types. Each point is one tumor sample, grey line is the best fit from linear model. Cor: Spearman correlation coefficient. Gene signature scores were adjusted by tumor content using linear regression. C. Association heatmap between one TE subfamily and multiple gene signatures and estimated immune infiltrates across 25 TCGA cancer types. Left: LTR21B. Right: MER57F. Color: Spearman correlation coefficient (cor) from partial correlation adjusting for tumor purity. Significance of correlation: * abs(cor)>0.5 & FDR<0.05, ** abs(cor)>0.5 & FDR<0.01; *** abs(cor)>0.5 and FDR<0.001. Bottom bars show the differential expression log2 fold change and FDR values of TE in each cancer type. Magenta: up-regulated. Green: downregulated. Grey: either no normal samples available or the TE expression level was too low for a given cancer type.

For type I IFN response expression, the mean r-squared values across cancer type was 0.67±0.21 in the full lasso model compared to 0.24±0.11 in the cellularity model. MamGypLTR2b (Gypsy), LTR21B (ERV1), L1PBb (L1) and MER57F (ERV1) were identified as TEs with the strongest positive association to type I IFN response (**Fig. 4B, Fig. 4C, Fig. S4D**). Type I IFN response is indicative of activation of cell-intrinsic anti-viral pathways and has been suggested to be induced by intracellular sensing of dsRNA formed from TE transcripts (*7, 8, 23*). These models suggest that total TE expression could be a major contributor to type I IFN activities in the tumor. Direct correlation analysis with estimated tumor immune infiltrates (*33*) also revealed for several cancer types positive association of LTR21B and MER57F to tumor plasma dendric cell (pDC) expression (**Fig. 4C**), which is consistent with known biological function of pDC as a potent producer of type I IFN.

Several HERV subfamilies, LTR21B, MER57F and HERVL74-int (ERVL) were also identified as the top TE correlates to expression levels of type II IFN response, CD8 T effector activity and immune checkpoint. Direct correlation analysis with estimated immune infiltrates (*33*) confirmed positive association of LTR21B and MER57F to CD8+ Tells expression (**Fig. 4C**).

Given the large explanatory power observed for DDR and immune response pathways, we next explored the directionality and strength of association from all individual TE subfamilies to these two biological systems using standard correlation analysis (**Fig. S4F**). We identified striking positive correlations with DDR from a large number of TE subfamilies in KIRC (a.k.a. renal clear cell carcinoma, 327 subfamilies), pancreatic adenocarcinoma (PAAD, 111 subfamilies) and sarcoma (SARC, 51 subfamilies), suggesting TEs may be a significant contributor to DDR for these 3 cancer types. KIRC has been previously observed as an immunogenic type of tumor that responds well to immunotherapy yet it tends to have low tumor mutation burden (TMB)(*15*). As the overall level of DDR related expression in KIRC is comparable to many cancer types (**Fig. S4G**), one possible explanation for the observed extensive positive correlations in KIRC is that TEs may be a significant contributor to DDR. Somatic TE transposition events have been previously described in TCGA samples to lead to insertional mutations private to tumors (*35*) and retrotransposition is known to create DNA double-strand breaks (*36*). KIRC was recently observed to harbor the highest proportion of insertion-and-deletion tumor mutations compared to other TCGA cancer types (*37*). DDR is known to activate immune signaling and inflammation (*38*). In light of our observation of extensive correlation between TE expression and DDR expression, more research is needed to elucidate the connection between TE expression, DDR and immunogenicity.

### 5-aza-2’-deoxycytidine treatment of glioblastoma cell lines induces TE expression and TE-derived peptide presentation on MHC class I molecules

While TE expression may contribute to innate immune activation and result in tumor inflammation, it may also contribute to the adaptive immune infiltration through presentation of TE-derived peptides on tumor cells (**Fig. 5A**). Certain HERV transcripts have been shown to result in MHC class I-bound peptides at tumor cell surface and serve as triggers for cytolytic T cell response (*10, 11*). We postulated that a variety TE peptides may be presented by tumor and subject to surveillance by the adaptive immune system.

**Fig. 5.**
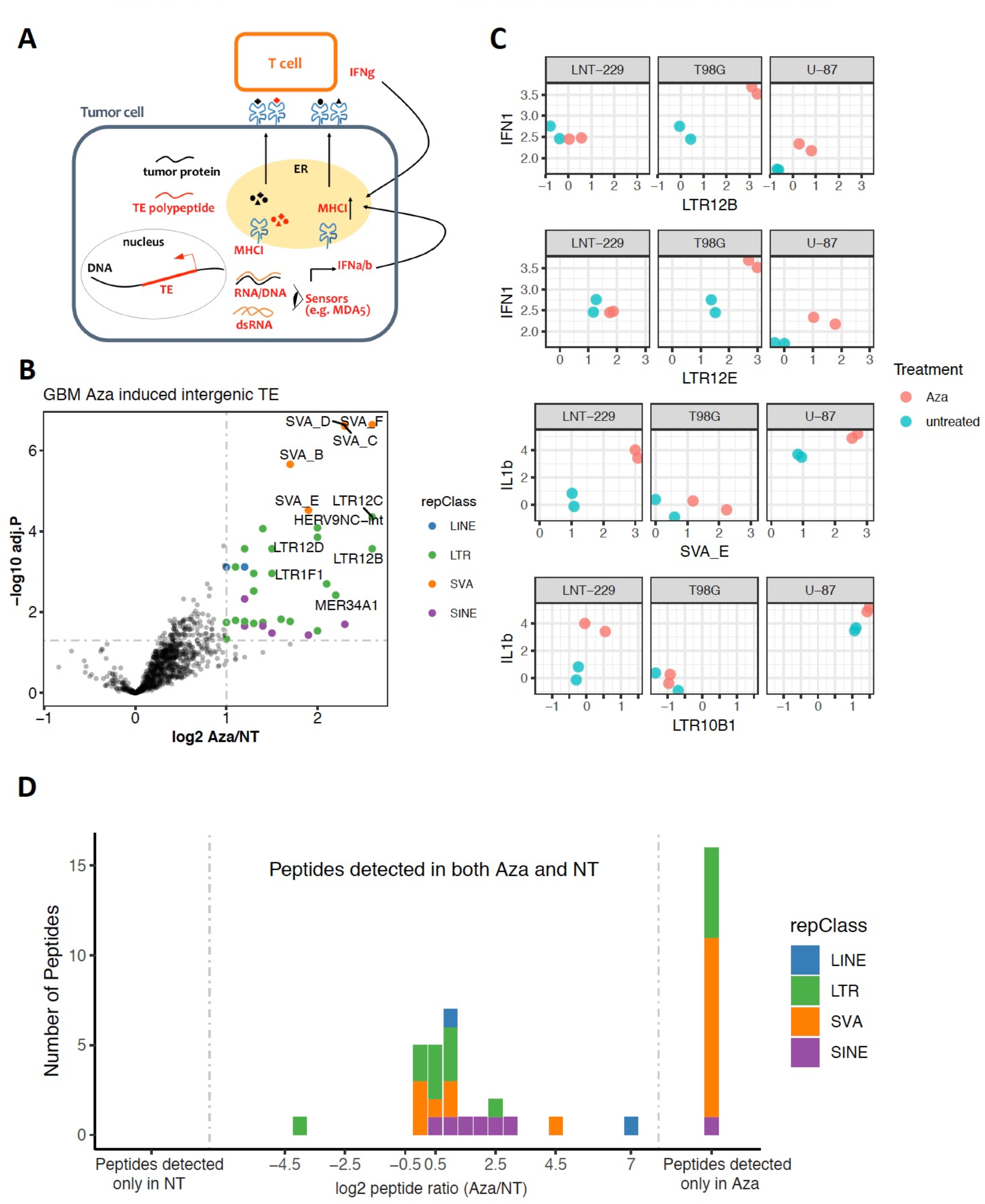
Decitabine treatment of GBM cell lines results in increased TE expression and TE-derived MHC-I peptide presentation. A. Working model of the impact of TE expression in the tumor. TE expression in the cytoplasm may trigger intracellular sensing of TE mRNA and result in type I IFN response. TE may be a source of tumor-associated antigens that can be presented at the tumor cell surface and recognized by TE-antigen specific T cells B. Volcano plot showing differential intergenic expression of TE subfamilies, Aza (decitabine) vs. NT (non-treated). TE subfamilies are colored by class at the significance threshold of log2FC > 1 and adjusted p-value < 0.05 and labeled if log2FC>1.5 and adjust p-value < 0.01. C. Association between select over-expressed TE subfamilies and cytokine gene signatures D. Effect of decitabine treatment on TE peptide presentation. Middle panel: histogram on log2FC for TE peptides abundance TE subfamilies with over-expression of mRNA. The log2FC of peptide presentation was calculated by comparing spectral areas for each peptide in both Aza vs. NT conditions. Peptides detected only in Aza: peptides uniquely detected in the treated condition. No TE peptides were detected only in the NT condition

To pursue this question, we re-analyzed a published multi-omic dataset (*39*) in which the authors originally examined the treatment effect of 5-aza-2’-deoxycytidine (decitabine, an inhibitor of DNA methyltransferase 1) on GBM cell lines through combined transcriptome, proteome and MHC class I peptidome analysis. Applying *REdiscoverTE* to the GBM transcriptome generated with rRNA depletion library prep, we discovered a much higher proportion of TE transcripts (~2.5%) and an enrichment of intronic TE expression in comparison to the data from poly-dT library preps, independent of treatment group (**Fig. S5A, Fig. S5B**). Epigenetic de-repression by decitabine resulted in strong over-expression of TEs originating from intergenic and intronic regions in these GBM cell lines, particularly TEs from SVA (SVA_B, C, D, E, F), ERV1 (n = 27), L1 (n=2) and Alu (n = 9) families (FDR < 0.05 and FC > 2, **Fig. 5B, Fig. S5C, Table S9**).

We searched the matching MHC class I peptidome and whole proteome data for translational products of TE by performing peptide identification and label-free quantification based on an augmented human proteome that included TE sequences from 51 over-expressed TE subfamilies. Using this approach, we identified 83 unique MHC-presented peptides derived from TEs and chose a subset of 39 peptides that were detected at least 3 times across all samples for further analysis. The majority of peptides mapped to TE elements resided in the intergenic regions of the reference genome, and some mapped to intronic regions (**Table S10**). These peptides were derived from over 10 subfamilies belonging to SVA (n=17), LTR (n=13), SINE (n=7) and LINE (n=2), with two subfamilies (SVA_D and LTR12C) representing half of the 39 peptides. Additionally, we identified 19 peptides derived from five of these subfamilies within the whole proteome data, adding further confidence in the translation potential of these TEs (**Table S11**). Sixteen (41%) peptides were detected only in the decitabine-treated condition, suggesting possible presentation of novel peptides, induced by a DNA demethylation agent. Seven of the remaining 24 peptides showed a two-fold increase in abundance under decitabine treatment compared to untreated condition (**Fig. 5C**). A subset of the peptides detected under both treated and untreated conditions were synthesized; we were able to verify their identity by mass spectrometry and confirm their binding to MHC-I molecule by peptide exchange assay (**Table S10**).

Simultaneous to TE expression increase, decitabine treatment also resulted in a striking induction of host gene expression, including many cancer testis antigen (CTA) genes such as the MAGE family genes, CYP1B1 and MELTF (**Fig. S5D**), as also reported in Shraibman et al. (*39*).

Geneset enrichment analysis confirmed that decitabine treatment is associated with the over-expression of not only CTA related pathways -- spermatogenesis (FDR = 5.1 x 10^-3^), allograft rejection (FDR = 1.3 x 10^-2^), but also a number of immune and cellular pathways that is consistent with above TCGA finding: inflammatory response (FDR = 4.0 x 10^-7^), TNFa signaling (FDR = 1.5 x 10^-3^), EMT (FDR = 6.4 x 10^-2^) and P53 response (FDR = 0.03, Table S12). Also consistent with above TCGA results are the observations that the expression of several TE subfamilies correlated with either IL1beta response or type I interferon response (**Fig. 5B**). Interestingly, expression of two subfamilies with the most number of peptides detected: SVA_D and LTR12C, were both strongly associated with DNA damage repair and homologous recombination in the TCGA KIRC cohort.

## Discussion

Our analysis revealed extensive TE expression in tumors, which strongly associated with the expression of innate immune genes and triggered the production of polypeptides that are processed and presented on MHC I molecules. These finding were made possible due to the development of a novel computational approach that simultaneously quantified the expression of host genes and all annotated repetitive elements. Key findings included the demonstration of transcriptional activities from all 5 classes of TE, including DNA transposons, with an enrichment seen in the evolutionarily youngest TEs. Much of the TE expression was derived from intergenic loci, supporting autonomous expression of many TEs rather than read-through resulting from host, protein-coding genes. Advancing prior observations of global demethylation in cancer, *REdiscoverTE* permitted the demonstration that TE expression is tightly linked to proximal DNA demethylation in tumors. Importantly, our results demonstrated that the expression of TEs not only is associated with increased innate immune responses but also results the presentation of TE-derived peptides on tumor cell surfaces by MHC-class I molecules for the engagement of adaptive immunity. Taken together, these findings suggest that TE expression in the tumor, whether spontaneous or induced under epigenetic therapy, has important clinical implications for cancer immunotherapy (*9, 40*).

Anti-viral response in the tumor has been demonstrated to potentiate patient response to immunotherapy (*7, 8*). In particular, for KIRC, where tumors have low mutational burden yet are highly immunogenic, HERV activity has been proposed to be a source of inflammation and mechanism of tumor regression(*15*). Analysis of point mutations alone, however, likely underestimates the true mutational burden (*37*). While the true extent of endogenously encoded antigens in the MHC-peptidome is not yet known, over the last 3 decades, about 10 HERV-derived peptides have been reported to be presented on a tumor cell surface, generate CD8+ clonal expansion and anti-tumor response in KIRC (*10, 11*), colorectal cancer (*41*) and melanoma (*42, 43*). Here, we demonstrate that LTR, LINE, SINE and SVA transcripts from intronic and intergenic regions could potentially be translated and presented on the MHC class I molecules of tumor cells.

Human DNA transposons are widely believed to be immobile (*23*), and their transcription has seldom been described in the literature. We have detected transcripts from a number of DNA transposons and identified MER75 as a top correlate to tumor DNA damage response. MER75, first reported in the original paper for the Human Genome Project, is a member of the piggyBac DNA transposon family and one of the youngest DNA transposons (*1*). piggyBac DNA transposons, in addition to being widely used as a genomic engineering tool to create insertional mutagenesis (*44*), are also ancestral to several domesticated host genes, such as PGBD5. PGBD5 was recently reported to encode an active DNA transposase expressed in the majority of childhood solid tumors and is responsible therein for site-specific oncogenic DNA rearrangements that require end-joining DNA repair (*45*). Together these findings suggest the possibility that activity of human DNA transposons may result in DNA damage, consistent with their known roles in chromosomal rearrangement and mutagenesis in other species (*46, 47*). However, it is still unclear whether TE activities by either DNA or re-retrotransposons, directly contribute or reflect other causes of genomic instability. Further experimental validation is necessary to establish the link between transcription of TEs such as MER75 and the tumor cellular DNA damage response. Nonetheless, in this study, we have provided strong evidence of correlation between TE expression and DDR in KIRC, pancreatic adenocarcinoma and sarcoma.

Regarding the computational methods presented herein, *REdiscoverTE* does not rely on consensus sequences, traditional short-read aligners, exclusion of multi-mapping reads, nor does it utilize step-wise operations that potentially introduce read-assignment bias (*13–16, 48*). In this study we have focused on TE expression in cancer, however, the method automatically quantifies expression of all repeats including those that are not transposable, e.g. satellites, which have been implicated in epithelial tumors (*49*). This method is broadly applicable to landscape profiling of the RE/TE expression in research areas beyond cancer, including autoimmune (*50*) and neurodegenerative diseases (*24, 51*), as well as normal embryonic stem cell development where TE activation is a hallmark of cellular identity and pluripotency (*13, 52–55*). Lastly, REdiscoverTE can be applied to transcriptomic analysis of TEs and REs for any organisms whose genome contain such elements (*12, 56, 57*).

In sum, our comprehensive in silico characterization of TE activities in tumors offers a number of predictions for experimental validation, and establishes a strong rationale for testing the newly identified class of TE-related, tumor-associated antigens as potential therapeutic vaccine targets.

## Acknowledgments

The authors are grateful to Avi Ashkenazi, Jieming Chen, Dorothee Nickles, Jeremie Decalf, Oliver Zill, Anna-Maria Herzner, Grace Xiao and Bob Yauch for their comments and feedback on the manuscript. We thank Jinfeng Liu for contribution to *REdiscoverTE,* Robert Piskol for support in pipeline development and William Forrest for statistical advice.

## Author contributions

Conceptualization: HC-H, SJ, MLA; Data curation: SL, PMH; Formal analysis: YK, HC-H, AC, CR; Funding acquisition: HC-H, JG; Investigation: HC-H, YK, AC, CR, MD; Methodology-REdiscoverTE: HC-H, SJ, YK, AC, RB; Methodology-TE peptides discovery-HC-H, CR, SJ; Methodology-peptide exchange: MD, CB; Project administration: HC-H; Resources: A-JT, CB; Software: YK, HC-H, AC, SJ, SL, PMH; Supervision: HC-H; Visualization: YK, AC, HC-H; Writing – original draft: HC-H, YK, AC, CR, MD, JG; Writing – review and editing: RB, SJ, CR, IM, MLA, JG, YK, HC-H.

## Data and materials availability

All data is available in the main text or the supplementary materials.

